# Deletion of Zyxin Reduces Endothelial Inflammation and Mitigates Atherosclerosis

**DOI:** 10.1101/2024.10.21.619550

**Authors:** Haozhong Huang, Yingzi Wang, Zhenyu Gao, Yi Feng, Tan Yang, Zihao Liu, Lei Li, Huimin Weng, Yuquan Tang, Linlu Liu, Yuhao Li, Zhixin Li, Yucheng Xie, Bin Liao, Fengxu Yu, Yongmei Nie

## Abstract

**BACKGROUND:** Oscillatory shear stress (OSS)-induced endothelial inflammation plays a critical role in the pathogenesis of atherosclerosis. However, the involvement of endothelial zyxin, a mechanosensor, in OSS-associated atherosclerosis and its underlying mechanisms remains unclear.

**METHODS:** To investigate the role of zyxin in vivo, Zyxin^iECKO^ApoE^−/−^ mice were utilized in atherosclerosis model induced by carotid artery ligation; in vitro, endothelial cells were subjected to disturbed flow in Ibidi systerm.

**RESULTS:** Zyxin was significantly upregulated in the OSS regions of both human and mouse arteries. The specific deletion of zyxin in endothelial cells (ECs) in ApoE^−/−^ (Zyxin^iECKO^ApoE^−/−^) mice reversed the ECs activation and atherosclerosis induced by OSS. In vitro studies indicated that the absence of zyxin reduced the induction of adhesion molecules and pro-inflammatory cytokines stimulated by OSS. Mechanistic investigations demonstrated that 14-3-3β facilitated yes-associated protein (YAP) phosphorylation at Serine 127, which played a critical role in retaining YAP within the cytoplasm. Under OSS stimulation, zyxin inhibited the phosphorylation of YAP at Serine 127 through its interaction with 14-3-3β, rather than direct regulation of YAP. This inhibition enhanced YAP’s nuclear translocation and promoted endothelial inflammation. Furthermore, it was shown that rosuvastatin inhibited zyxin expression in human umbilical vein endothelial cells and the vascular endothelium of ApoE^−/−^ mice. This inhibition led to decreased levels of inflammatory markers and adhesion molecules associated with atherosclerotic lesions observed in the partially ligated left common carotid arteries of ApoE^−/−^ mice.

**CONCLUSIONS:** This study further confirms that zyxin is a mechanoreceptor in endothelial cells and elucidates the indispensable role of the zyxin-14-3-3β-YAP axis in endothelial inflammation and atherogenesis. This indicates that Zyxin plays a crucial role in protecting against atherosclerosis.

## INTRODUCTION

Atherosclerosis (AS) serves as a significant pathological foundation for a variety of cardiovascular diseases (CVDs), including aortic aneurysms and coronary heart disease, posing a substantial threat to human health^1–3^. Notably, atherosclerotic plaques are predominantly found at arterial bends and bifurcations where characterized by low or disturbed shear stress—while their occurrence is markedly rare in straight arterial segments where laminar flow prevails^4–9^. Understanding the mechanisms driving the site-specific distribution of these plaques is essential for developing effective therapeutic strategies for AS.

As an initial pathogenic event in atherosclerosis, disturbed flow alters the morphology and cytoskeleton of ECs and modulates their intracellular biochemical signalling and gene expression, resulting in phenotypic and functional changes in ECs^5,10–12^. Hemodynamic signals continuously act on the vascular endothelial cells (VECs) lining the arterial wall, leading to the activation of numerous mechanosensors and responsive microdomains^4,12,13^. These mechanosensors detect biomechanical stimuli and transduce them into biochemical signals, thereby regulating endothelial cell homeostasis and vascular function in both healthy and diseased states^12,13^. Emerging evidence highlights the activation of various mechanosensory molecules and complexes—such as intercellular junction proteins, adhesion molecules, membrane microdomains, ion channels, and glycocalyx—by both laminar and disturbed flows, resulting in atheroprotective or atherogenic phenotypes^4,5,12,13^. This modulation further influences intracellular biochemical signals and gene expression^5,10,12^, with the inflammatory responses induced by these cascades being critical to the mechanisms underlying AS^5,10,14^, although the precise pathways involved remain to be elucidated. The mechanosensitive protein zyxin has been highlighted in various studies for its dual role in cellular mechanics^15^. It translocates biochemical signals to the nucleus in response to mechanical forces, regulating gene expression, or localizes along stress fibers to facilitate actin polymerization^15,16^. Primarily found at cell-extracellular matrix (ECM) and cell-cell junctions, zyxin is associated with actin stress fibers and focal adhesion plaques (FAs)^17^. As a member of the LIM (Lin11, Isl-1, and Mec-3) domain family, zyxin is highly conserved and crucial for mechanotransduction^18,19^. FAs, located at the ends of actin filaments, serve as sites for force transmission^19^. Zyxin comprises two distinct motifs: an N-terminal proline-rich domain that interacts with partners like α-actinin and a C-terminal LIM domain essential for its force-sensing function^18–20^. Recent research underscores zyxin’s importance in cardiovascular health, mediating endothelial cell responses to mechanical stretching and influencing inflammatory gene expression^21–23^. Loss of zyxin function promotes a synthetic phenotype in vascular smooth muscle cells, contributing to aortic aneurysm development^23^. In zyxin-deficient mouse models, increased vascular pressure worsens myocardial apoptosis and heart failure symptoms^23^. Nevertheless, the specific mechanism by which endothelial zyxin operates in AS within the OSS microenvironment remains to be explored. Atheroprone disturbed flow has been demonstrated to activate YAP, a downstream effector of the Hippo pathway, promoting inflammation and atherogenesis^24–26^. An unresolved question in YAP signaling is how external mechanical cues are transduced to regulate YAP^24,26,27^. Previous studies suggest that the 14-3-3 protein regulates YAP nuclear translocation by modulating its phosphorylation state^28,29^. Additionally, zyxin has been implicated in breast cancer development by regulating YAP activity in tumor cells^30^. Dynamic stretching instigates the activation of zyxin and expedites the translocation of YAP into the nucleus; conversely, loss of zyxin function hinders this process^21,22^. This implies that mechanical stimulation may induce YAP nuclear translocation through a zyxin-mediated pathway^21,22,30–32^. However, the mechanisms by which endothelial zyxin senses flow signals and coordinates YAP remain poorly understood.

This study demonstrates that zyxin functions as a mechanosensor in ECs, regulating cellular responses to shear stress and influencing the site-specific distribution of AS. We provide evidence for the role and mechanisms by which zyxin regulates YAP nuclear translocation in response to OSS stimulation. Our findings indicate that 14-3-3-β, a crucial member of the 14-3-3 protein family, contributes to stabilizing YAP phosphorylation at Serine 127 and retaining it in the cytoplasm. OSS-activated zyxin participates in the pathogenesis of AS by interacting with the 14-3-3-β protein (YWHAB) to facilitate YAP nuclear translocation and enhance the expression of downstream inflammatory genes.

## METHODS

### Data Availability

All supporting data are included in the article and its Supplemental Materials. For details on the experimental procedures and materials used, please refer to the Major Resources Table and the Materials and Methods section in the Supplemental Materials.

## RESULTS

### 1. Zyxin upregulated in human and mouse atherosclerotic plaques

To evaluate the expression of zyxin in atherosclerotic plaques, samples of non-atherosclerotic and atherosclerotic plaques were collected from patients undergoing coronary bypass surgery at the Department of Department of Cardiovascular Surgery, Affiliated Hospital of Southwest Medical University. This study was conducted in accordance with established inclusion and exclusion criteria (refer to Table S1 for patient clinical information). The collected arterial tissue samples underwent H&E staining, and representative images of non-atherosclerotic and atherosclerotic plaques in human arterial tissue are presented in Figures S1A-B. RT-qPCR (real-time polymerase chain reaction) analysis demonstrated that zyxin mRNA transcripts were significantly elevated in human atherosclerotic plaques compared to non-atherosclerotic tissues (Figure 1A). Consistent with the mRNA data, Western blot analysis indicated that zyxin protein levels were also significantly higher in atherosclerotic plaques (Figure 1B). Given the heterogeneity of cell types in human arterial tissues, an immunofluorescence approach was employed to examine zyxin expression specifically in ECs. The results revealed that zyxin fluorescence intensity was markedly greater in atherosclerotic plaques compared to non-atherosclerotic arterial tissues (Figure 1C). To investigate the effect of hemodynamics on zyxin expression in vivo, thoracic aorta and aortic arch tissues were harvested from ApoE^−/−^ mice fed a high-fat diet and stained with oil red O to identify atherosclerotic lesions. A substantial increase in the area of oil red O staining was observed in the aortic arch (medial surface) compared to the thoracic aorta (Figure 1D). RT-qPCR analysis further indicated that zyxin mRNA transcripts were expressed at higher levels in the aortic arch than in the thoracic aorta (Figure 1E). Moreover, the levels and nuclear localization of zyxin protein were examined in various regions of the mouse aorta. In the aortic arch of ApoE^−/−^ mice, zyxin protein levels were elevated compared to those in the thoracic aorta (Figure 1F). Immunofluorescence analysis of ApoE^−/−^ mouse vascular tissue further revealed that zyxin fluorescence intensity was significantly greater in the aortic arch than in the thoracic aorta (Figure 1G). Notably, there was no significant difference in zyxin levels within the nuclei at the inner curvature of either the thoracic aorta or the aortic arch (Figure 1G, Figure S1C). These results suggest that endothelial zyxin may play a role in the progression of AS.

**Figure 1.**
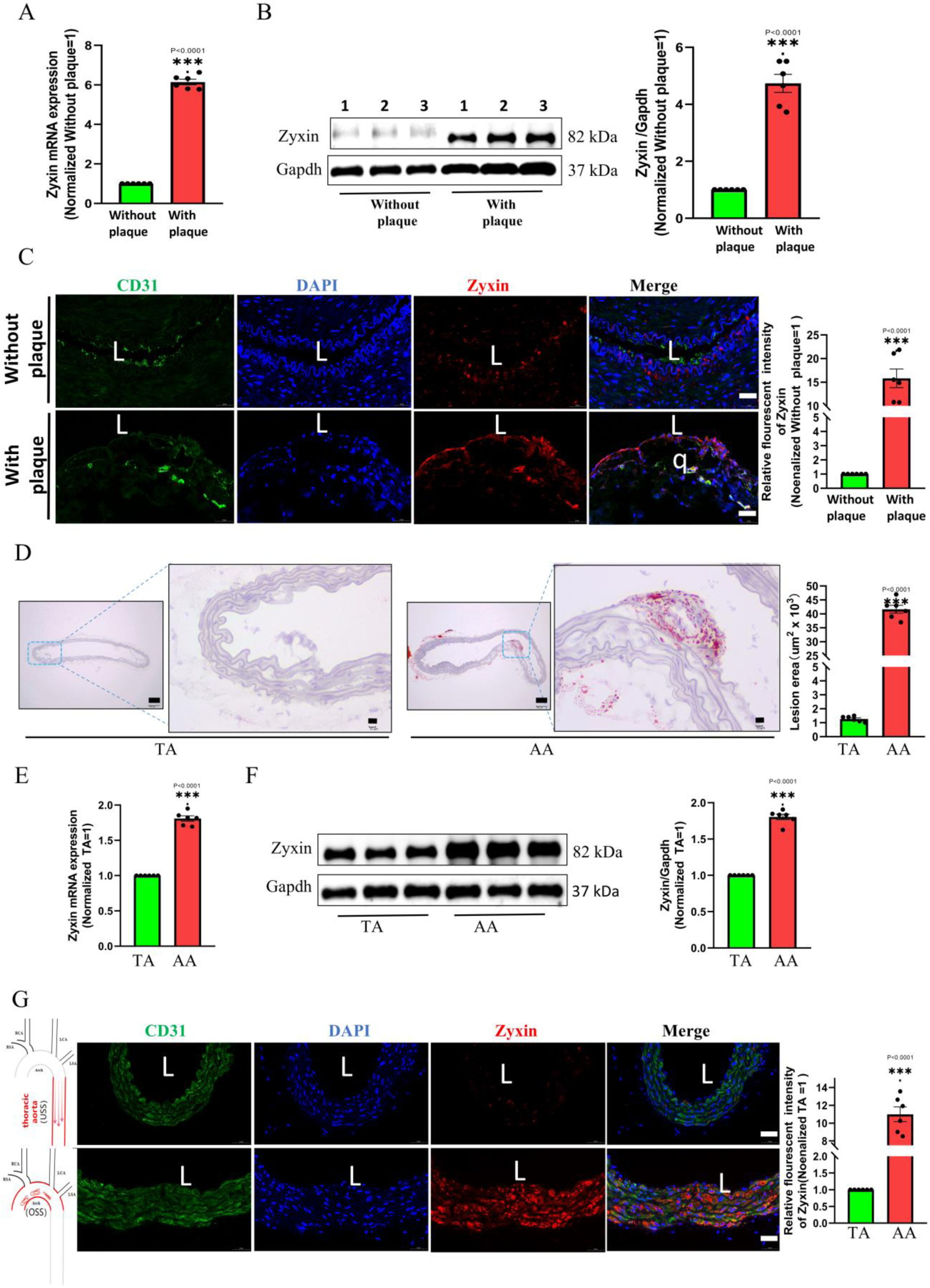
Zyxin Is Upregulated In Human and Mouse Atherosclerotic Plaques. A. Zyxin mRNA expression was significantly upregulated when comparing human plaque groups with matched non-plaque tissues (n = 6). **B.** Representative and summarized Western blotting data demonstrating the expression of zyxin protein level in human arterial tissue. zyxin protein expression was elevated in atherosclerotic plaques in contrast to non-plaque arterial tissues (n = 6). **C.** Immunofluorescence staining for zyxin ^32^, CD31 ^54^ and DAPI (blue) in human arterial tissue. Human arterial tissues were classified into without-plaque and with-plaque groups. Scale bars: 50 μm, L, lumen; P, plaque. (n = 6). **D.** Representative images of oil red O staining of the thoracic aorta and rat aortic arch in high-fat diet ApoE knockout (ApoE^KO^) mice and quantification of the lesion area (n = 6). Zyxin was increased in the aortic arch (AA, curved, the lower panel) compared to the thoracic aorta (TA, straight, the upper panel). Scale bars: Original image 100 μm, enlarged image 10 μm. **E.** zyxin mRNA expression was significantly upregulated when comparing human plaque groups with matched non-plaque tissues (n = 6). **F.** Immunoblotting showing that the zyxin protein level was higher in the AA compared with the TA from high-fat diet ApoE^KO^, with quantification of protein expression in the right panel (n = 6). **G.** En face immunofluorescence staining of zyxin, CD31^32,54^, and DAPI (blue) in the ApoE^-/-^ mouse aorta, showing an increased fluorescence intensity of zyxin in the inner curvature of the AA compared with the TA. The quantification data were presented in the right panel (n = 6). Scale bars: 50 μm, L, lumen; P, plaque. Statistical analysis was performed by nonparametric Mann-Whitney test for A-G.

### 2. Zyxin prominently implicated in atherosclerosis due to OSS

Numerous studies demonstrated that atherosclerotic plaques tend to form in regions subjected to OSS. To assess the effects of different shear stress modalities on zyxin expression levels, human umbilical vein endothelial cells (HUVECs) were exposed to OSS or uniform shear stress (USS) generated by the Ibidi pump system. Western blot assays indicated that zyxin protein expression increased in response to OSS compared to static cells (STA), with the most significant upregulation observed after 6 hours of OSS stimulation (Figure S2A). Vascular cell adhesion molecule-1(VCAM-1) expression levels were subsequently examined under various shear stress conditions, yielding similar results through Western blot analysis (Figure S2B). RT-qPCR analysis of HUVECs and human aortic endothelial cells (HAECs) exposed to OSS for 6 hours revealed that OSS enhanced the mRNA expression of zyxin, VCAM-1, and Intercellular cell adhesion molecule-1 (ICAM-1) in both cell types (Figure 2A-B, Figure S2C-D). Additionally, zyxin expression and the protein levels of the inflammatory factor VCAM-1 were upregulated by OSS stimulation (Figure 2C, Figure S2E). To delineate the role of zyxin in ECs activation and AS resulting from disturbed flow in vivo, partial ligation of the left common carotid artery was performed in ApoE^−/−^ mice (illustrated in Figure 2D). A Western diet was administered immediately following surgery to accelerate the development of AS^29,33^. Carotid ultrasonography was conducted weekly for four consecutive weeks postoperatively to confirm disturbed flow in the ligated left carotid artery (LCA) and to compare it with the control group (Figure 2E, Figure S3A). Eight-week-old male ApoE^−/−^ mice underwent either sham surgery or partial ligation of the left common carotid artery. Four weeks after the surgical intervention, the left common carotid arteries of both sham-operated and experimental cohorts were harvested for Oil Red O and H&E staining. A significant increase in the area of atherosclerotic lesions was observed in ApoE^−/−^ mice in the PLCA group compared to the sham group (Figure 2F, Figure S3C). Immunofluorescence labeling results showed that the fluorescence intensity of zyxin and the inflammatory factor VCAM-1 was significantly greater in the surgical group compared to the sham-operated group (Figure 2G), supported by Western blot and RT-qPCR assays (Figure S3D-J). In contrast, no significant differences were observed in serum triglyceride (TG), total cholesterol (CHO), low-density lipoprotein (LDL-C), or high-density lipoprotein (HDL-C) levels in ApoE^−/−^ mice on a high-fat diet (Figure S3K-N). These findings indicate that OSS upregulates zyxin expression and enhances the levels of inflammatory factors in ECs involved in atherogenesis.

**Figure 2.**
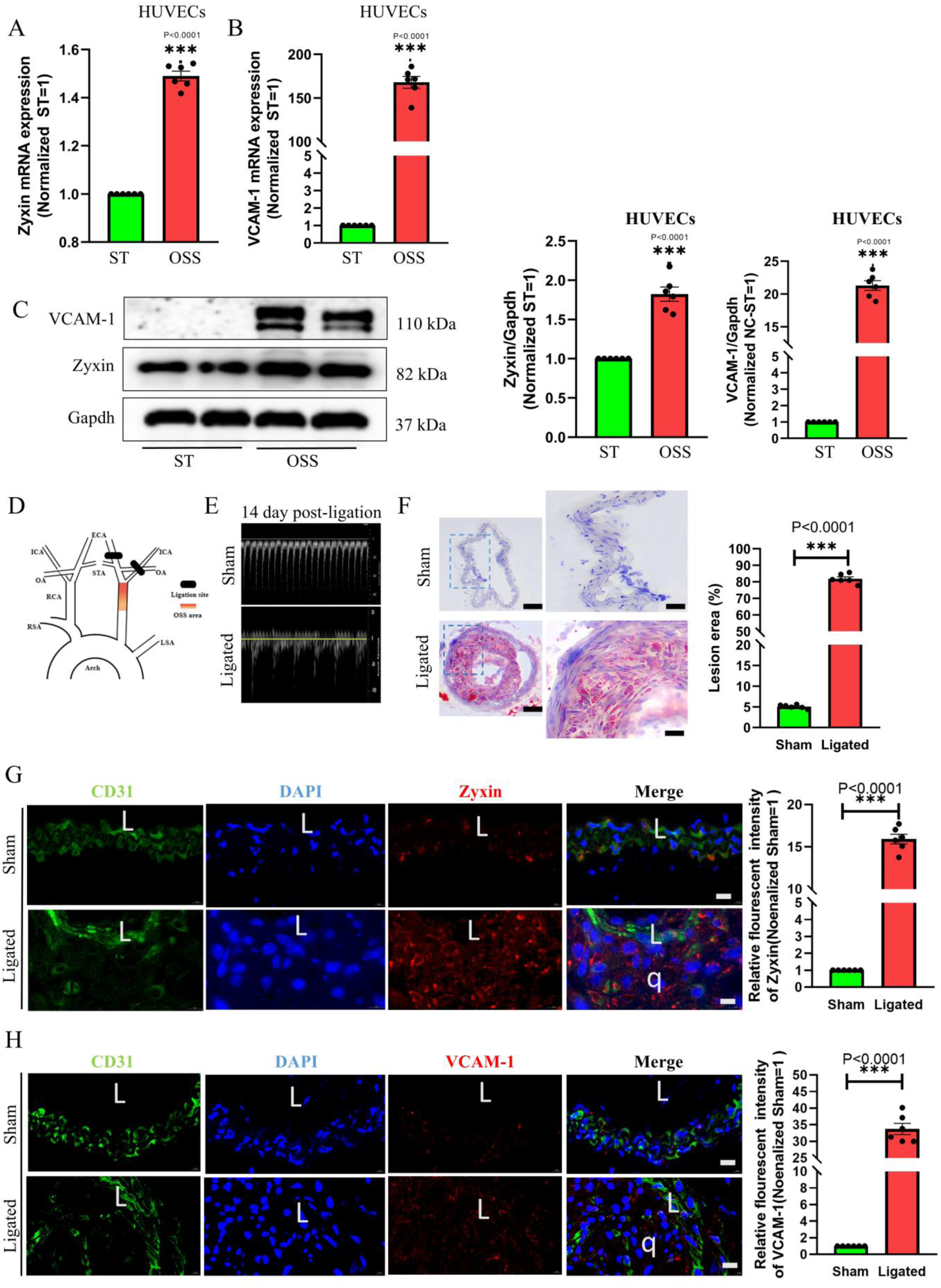
Zyxin is implicated in OSS-related atherosclerosis. **A-B,** Oscillatory shear stress (OSS) for 6 h upregulated zyxin (A) and VCAM-1 (B) mRNA levels in human umbilical vein ECs (HUVECs), n=6. **C,** Representative and summarized western blot data showing inhibited zyxin and VCAM-1 protein expression in HUVECs exposed to OSS for 6 h, n=6. **D,** Diagram of the partial ligation model, black shading indicates the ligation site and red shading indicates the OSS area. **E,** Ultrasound images showing flow velocity profiles and revealing that partial ligation induces flow reversal in the ligated left carotid artery (LCA) during diastole. Representative images were obtained on days 0 and 14 after surgery. **F,** Arterial tissues were isolated to examine atherosclerotic lesions after 4 weeks of ligation, Ligated LCA was sectioned for Oil Red O staining and quantification of the lesion area (n=6). Scale bar: Original image100μm, enlarged image 10μm. **G-H,** LCA was isolated for en face immunostaining of Zyxin (G) and VCAM-1 (H) in ECs 4 weeks after ligation. Quantification of relative fluorescence intensity (right). Green, CD31; Blue, DAPI, red, zyxin or VCAM-1. L, lumen; q, plaque. n = 6. Scale bar: 10 μm. Statistical analysis was performed by nonparametric Mann-Whitney test for A-C and F-H.

### 3. Specific knockdown of zyxin in vascular endothelial cells reduced atherosclerotic lesions in ApoE−/− mice

The crucial role of zyxin in modulating pro-inflammatory gene expression leads us to hypothesize that endothelial zyxin significantly contributes to the pathogenesis of AS, a chronic inflammatory disorder affecting blood vessels. To investigate this hypothesis, we examined the role of zyxin in the progression of AS in ApoE^−/−^ mice by administering adeno-associated virus ZYX (AAVENT-Zyx-RNAi) to induce a specific knockout of endothelial zyxin in these mice, designated as Zyxin^iECKO^ ApoE^−/−^. Following screening, the optimal knockdown site among three candidates was identified (Figure S4A-C). The selected construct or saline was then injected into ApoE^−/−^ mice in the Zyxin^iECKO^ and Zyxin^WT^ groups, respectively, via the tail vein. Seventy-two hours later, the mice in each group underwent either sham surgery or partial ligation of the left common carotid artery (PLCA). Postoperatively, all ApoE^−/−^ mice were placed on a high-fat diet. Four weeks after surgery, intact aortas—including bilateral common carotid and iliac arteries—were isolated and harvested from each group. Zyxin deletion in ECs reduced both plaque area and Oil Red O staining in the en face aortas of male mice compared to wild-type (WT) mice (Figures 3A and 3B). Histological analysis of the left common carotid artery indicated that the absence of endothelial zyxin was associated with a reduction in atherosclerotic plaque size (Figure 3C). Additionally, the levels of the inflammatory protein VCAM-1 were significantly lower in the left common carotid artery of Zyxin^iECKO^ mice compared to WT mice (Figure 3D). Collectively, these data suggest that zyxin-specific deletion in vascular ECs effectively reduces the size of atherosclerotic lesions in ApoE^−/−^ mice.

**Figure 3.**
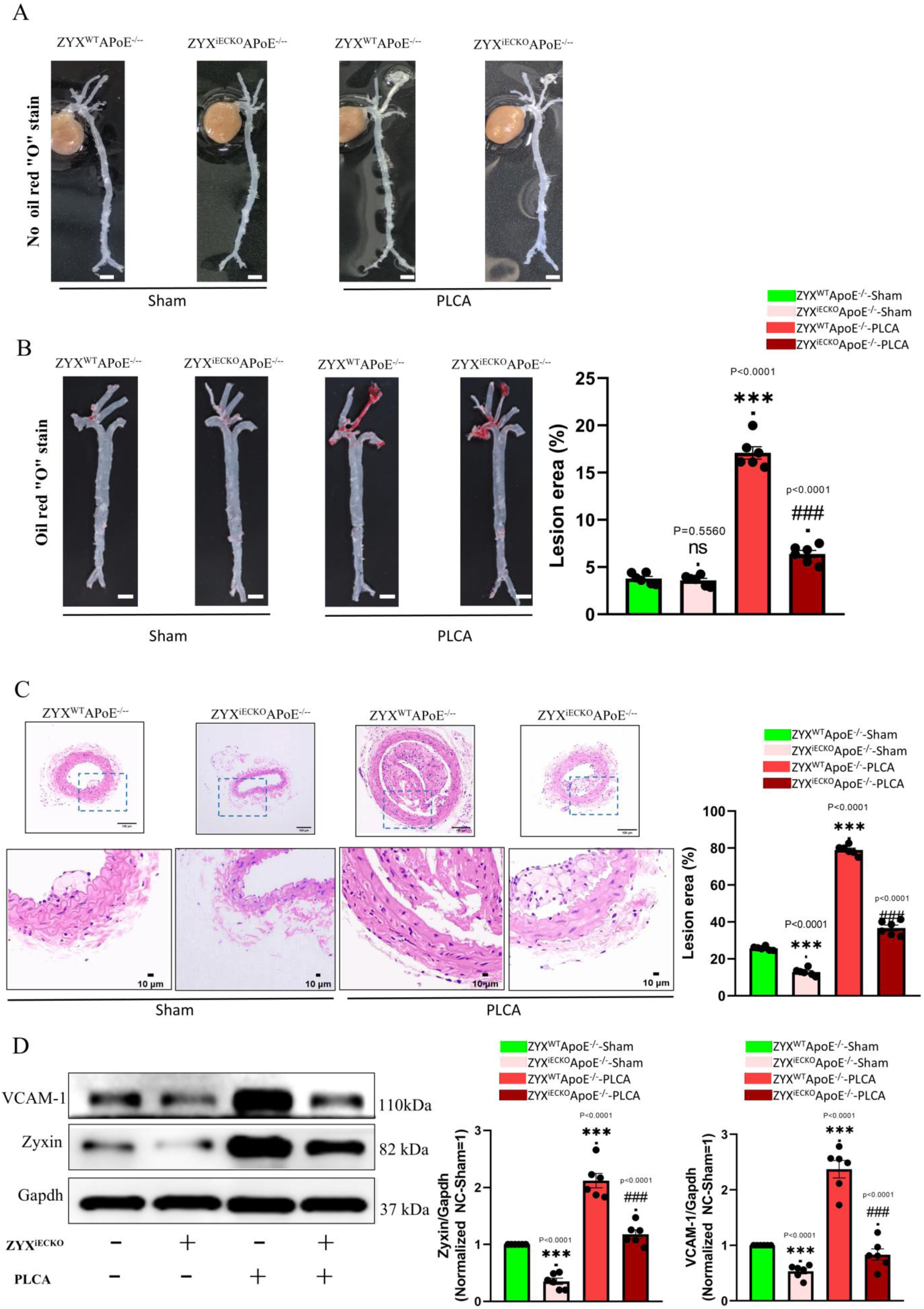
**A–C,** Arterial tissues were isolated to examine atherosclerotic lesions 4 weeks after ligation. **A,** No oil red staining; scale bar: Scale bar: 1mm, n=6. **B,** Arterial tissues were used for oil red staining and quantification of the lesion area. Scale bar: 1mm, n=6. **C,** Ligated LCA was sectioned for hematoxylin and eosin staining and quantification of the lesion area (n=6). Scale bar: Original image 100μm, enlarged image10μm. L, lumen; P, plaque. **D,** Immunoblotting showing zyxin and VCAM-1 protein levels in the left common carotid artery after sham operation or ligation of Zyxin^WT^ and Zyxin^iECKO^ mice. Quantitative data are shown in the figure ring, n=6. Statistical analysis was performed by 2-way ANOVA followed by the Tukey test for B-D.

### 4. Zyxin deficiency in endothelial cells inhibits the expression of inflammatory factors and adhesion molecules

Based on the inflammation viewpoint of AS, The impact of endothelial zyxin deficiency on ECs inflammation was subsequently investigated. Notably, a significantly elevated fluorescence intensity of CD31-labeled VCAM-1 was observed in ECs within atherosclerotic plaques when compared to non-plaque arterial tissue in humans (Figure S5A). In Zyxin^iECKO^ApoE^−/−^ mice, the specific depletion of endothelial zyxin resulted in a reduction of VCAM-1 fluorescence intensity in the aortic arch vascular tissue following partial ligation of the left carotid artery (Figure 4A). To explore the precise mechanisms by which zyxin mediates atherogenesis, we first conducted and verified the deletion of zyxin function in HUVECs (Figure S5B-C), examining the expression of adhesion molecules, pro-inflammatory cytokines, and chemokines. The downregulation of zyxin was found to reduce the expression of adhesion molecules (ICAM-1 and VCAM-1) and Monocyte chemoattractant protein-1 (MCP-1) in OSS-induced HUVECs (Figures 4B-F). These findings were confirmed by RT-qPCR and Western blot analyses and further verified in HAECs (Figure 4G). Consistent with these results, endothelial zyxin deficiency also inhibited the expression of TNF-α-induced adhesion molecules (ICAM-1 and VCAM-1) and the chemokine MCP-1 (Figure S5D-H). Collectively, these results suggest that endothelial zyxin contributes to atherogenesis by enhancing the expression of adhesion molecules and pro-inflammatory cytokines in ECs.

**Figure 4.**
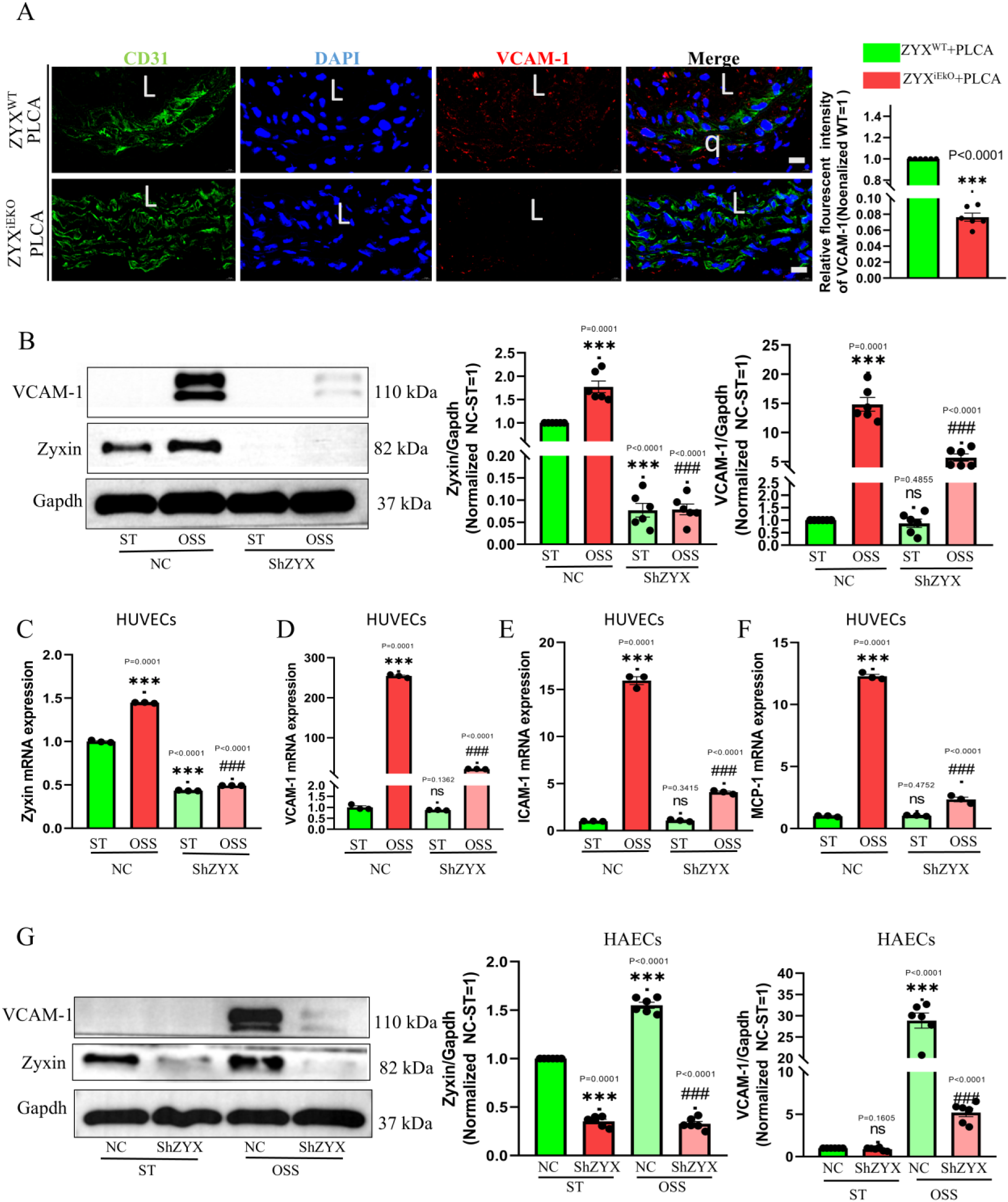
Zyxin deficiency in endothelial cells inhibits the expression of inflammatory factors and adhesion molecules. **A,** Representative images of immunostaining of VCAM-1 (vascular cell adhesion molecule-1) in LCA from male Zyxin^WT^ mice and male Zyxin^iECKO^ mice following 4 weeks of ligation, showing decreased expression of inflammatory molecules in Zyxin^iECKO^ mice, with quantification data in the right panel (n=6 mice per group). VCAM-1 ^32^, CD31 ^54^, and DAPI (blue) staining. L, lumen; q, plaque. n = 6, Scale bar: 10 μm. B, Immunoblotting showed that the protein levels of VCAM1 were decreased in HUVECs transfected with Zyxin shRNA compared with those transfected with Con shRNA, with or without OSS treatment. The quantification data are shown in the lower panel, n=6. **C-F,** zyxin, VCAM-1, ICAM-1 and MCP-1 mRNA expression was decreased in HUVECs transfected with Zyxin shRNA compared with those transfected with Con shRNA, with or without OSS treatment for 6 h, n=3. **G,** Immunoblotting showed that VCAM-1 protein levels were decreased in HAECs from zyxin shRNA compared with those of Con shRNA, with or without OSS treatment. The quantification data are shown in the lower panel, n=6. Statistical analysis was performed by 2-way ANOVA followed by nonparametric Mann-Whitney test for A, by 2-way ANOVA followed by the Tukey test for B-G.

### 5. Zyxin mediates the nuclear localization of YAP and promotes the expression of inflammatory factors in endothelial cells

We explored the potential mechanisms underlying zyxin-induced ECs inflammation through comprehensive transcriptome analysis of RNA sequencing from HUVECs subjected to different patterns of shear stress. KEGG pathway enrichment analysis revealed several enriched pathways crucial for cellular metabolism and proliferation, including DNA replication, the Hippo signaling pathway, the cell cycle, and the PI3K-Akt signaling pathways (Figure 5A). To determine whether zyxin interacts with key molecules within these signaling pathways, we examined the protein interaction network using STRING analysis. This revealed that YAP, a pivotal molecule in the Hippo signaling pathway, is potentially associated with zyxin (Figure 5B). To elucidate whether the OSS-induced increase in YAP nuclear localization is linked to zyxin, we conducted both loss-of-function and gain-of-function experiments in ECs using lentiviral vectors. The observed trends in YAP changes induced by zyxin loss-of-function or gain-of-function were consistent with those of zyxin at both the mRNA and protein levels (Figure S6A-E). Interestingly, endothelial zyxin deficiency inhibited the OSS-induced expression of zyxin, YAP, and its downstream target genes, including CTGF and Cyr61, as well as inflammatory factors such as VCAM-1, ICAM-1, and monocyte chemotactic protein-1 (MCP-1). These results were consistently verified at both the mRNA and protein levels (Figure 5C, Figure S6F-G). Furthermore, when HUVECs were pretreated with Verteporfin, a non-specific YAP inhibitor, for 30 minutes prior to static or OSS treatment for 6 hours, Verteporfin significantly downregulated YAP expression and inhibited the elevated expression of zyxin, YAP, and its downstream target genes, CTGF, Cyr61, along with inflammatory factors VCAM-1, ICAM-1, and MCP-1. Notably, the inhibitory effect of Verteporfin on YAP was more pronounced under OSS stimulation compared to static conditions (Figure 5D, Figure S6F-H). To further validate the effect of zyxin-mediated OSS on YAP nuclear translocation, we conducted cellular assays targeting the nucleoplasmic separation of YAP. We found that the increase in YAP due to OSS primarily manifested as an increase in the level of cytosolic YAP protein, with no statistically significant difference in YAP protein expression in the cytoplasm (Figure 5E). However, the protein level of cytosolic YAP was significantly reduced by the deletion of zyxin (Figure 5F). These findings suggest that the OSS-induced nuclear localization of YAP is mediated by zyxin, which enhances the expression of inflammatory factors in ECs.

**Figure 5.**
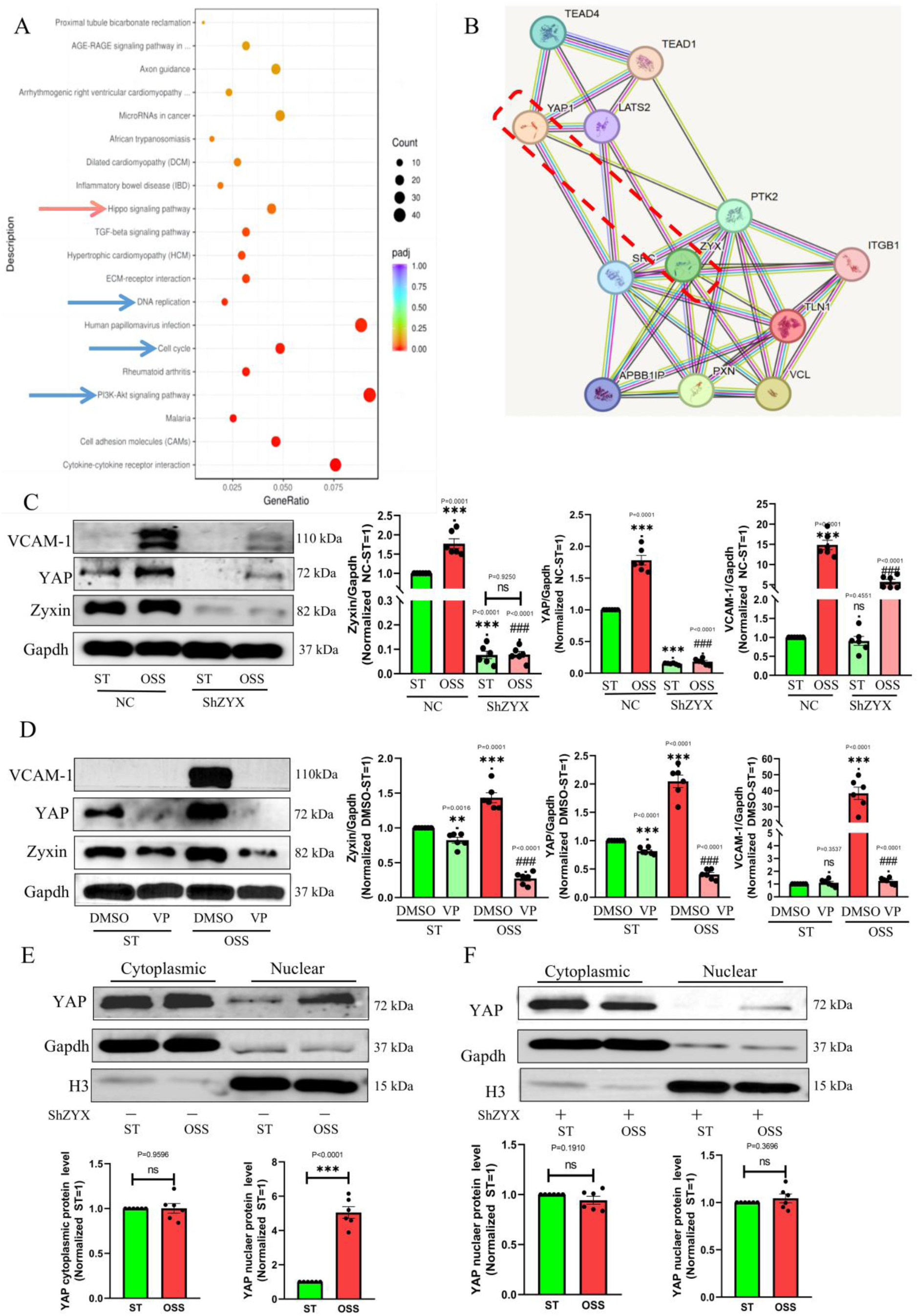
Zyxin mediates the nuclear localisation of YAP and promotes the expression of inflammatory factors in endothelial cells. **A,** HUVECs were exposed or not exposed to OSS for 6 h. RNA sequencing was performed for transcriptomic analysis of the KEGG pathway enrichment bubble map. **B,** The string protein interaction network shows the association diagram between zyxin and YAP, a key molecule of the Hippo signalling pathway. **C,** Immunoblotting showed that the protein levels of YAP and VCAM-1 were decreased in HUVECs transfected with zyxin shRNA compared with those transfected with Con shRNA, with or without OSS treatment. The quantification data are shown in the lower panel, n=6. ST, static; OSS, oscillatory shear stress. **D,** Immunoblotting showed that the protein levels of YAP and VCAM-1 were decreased in HUVECs from VP compared to those from DMSO, with or without OSS treatment. The quantified data are shown in the lower panel, n=6. ST, static; OSS, oscillatory shear stress. DMSO, Dimethyl sulfoxide ; VP, Verteporfin. E-F, Western blotting detected YAP protein levels in the cytoplasm or nucleus of HUVECs. HUVECs exposed to OSS for 6 hours resulted in increased YAP protein levels in the nucleus(E), which was reversed after zyxin loss-of-function(F), while there was no significant difference in cytoplasmic YAP protein levels. n=6, ST, static; OSS, oscillatory shear stress. Statistical analysis was performed by 2-way ANOVA followed by the Tukey test for C and D, by nonparametric Mann-Whitney test for E-F.

### 6. The zyxin-14-3-3 axis mediates YAP nuclear translocation to promote endothelial inflammation

The mechanisms by which zyxin regulates YAP nuclear translocation were explored. Initially, an effort was made to determine whether zyxin interacts with p-YAP(s127); however, results from the Co-IP experiment indicated that no direct interaction exists (Figure S7A). Building on previous work by our team, it was established that 14-3-3 is essential for retaining p-YAP(s127) in the cytoplasm and facilitating its transport into the nucleus. Additionally, the regulation of YAP signaling by OSS through the zyxin-14-3-3 axis was investigated. The 14-3-3 protein family currently comprises seven subtypes, and understanding their variability in response to OSS is crucial for subsequent experiments. The mRNA expression of each isoform within the 14-3-3 protein family in HUVECs was assessed using RT-qPCR under physiological conditions, OSS stimulation, and treatment with TNF-α (10 ng/ml). The results indicated that the SFN (σ) subtype was expressed at very low levels in HUVECs (Figure S7B-D). While YWHAE (ε) was identified as the subtype with the highest physiological expression (Figure S7B-D), the expression of 14-3-3-β (YWHAB) mRNA was most pronounced under OSS or TNF-α (10 ng/ml) treatment. Given the disparities in 14-3-3-β expression, HUVECs were transfected with lentivirus targeting the 14-3-3-β (YWHAB) subtype, and the mRNA expression levels of each isoform within the 14-3-3 family were assessed. The RT-qPCR results indicated that, with the exception of SFN (σ) isomers, the downregulation of 14-3-3-β (YWHAB) mRNA levels led to an upregulation of the mRNA levels of the other five isoforms compared to the negative control (NC) group. However, this did not reverse the reduction in total 14-3-3 protein expression caused by the deletion of 14-3-3-β (Figure S7E-F). Therefore, it was hypothesized that 14-3-3-β may be the predominant subtype of the 14-3-3 protein family responsive to OSS in HUVECs. Additionally, upon the transfection of HUVECs with lentiviruses for zyxin knockdown, a significant reduction in zyxin protein expression was observed, along with a marked decrease in total 14-3-3 protein levels in the zyxin knockdown group compared to the negative control (NC) group (Figure 6A). In instances where Sh-14-3-3β was knocked down via lentivirus transfection, both 14-3-3β and total 14-3-3 protein expressions were significantly diminished (Figure S7F). In contrast, no statistically significant difference in zyxin protein levels was shown compared to the NC group (Figure 6A). These findings suggest a strong association between zyxin and 14-3-3, with zyxin likely being positioned upstream to regulate 14-3-3 activity. To investigate whether the zyxin-14-3-3β axis modulates YAP signaling, HUVECs were exposed to OSS for 6 hours. This treatment resulted in the promotion of dephosphorylation of YAP at the Serine 127 (S127) locus and an increase in YAP expression (Figure 6B). However, in the absence of zyxin, total YAP protein levels were found to decrease, and phosphorylation at the S127 site was enhanced. Conversely, when 14-3-3β was knocked down, YAP phosphorylation at S127 was inhibited while total YAP expression was increased (Figure 6C). Subsequently, HUVECs were treated with 10 ng/ml of TNF-α or phosphate-buffered saline (PBS) for varying durations, and changes in zyxin and 14-3-3 protein expressions were found to be closely aligned (Figure S7G). Given that 14-3-3β may be a principal member of the 14-3-3 protein family, it was hypothesized that an interaction between zyxin and 14-3-3β occurs, as indicated by immunoprecipitation assays (Figure 6D). The accumulated evidence supports the notion that the 14-3-3β isoform is crucial for the stabilization of YAP retention in the cytoplasm, inhibiting its nuclear translocation. Consequently, it is concluded that zyxin regulates OSS-induced YAP nuclear translocation via 14-3-3β (YWHAB), thereby promoting inflammation in ECs.

**Figure 6.**
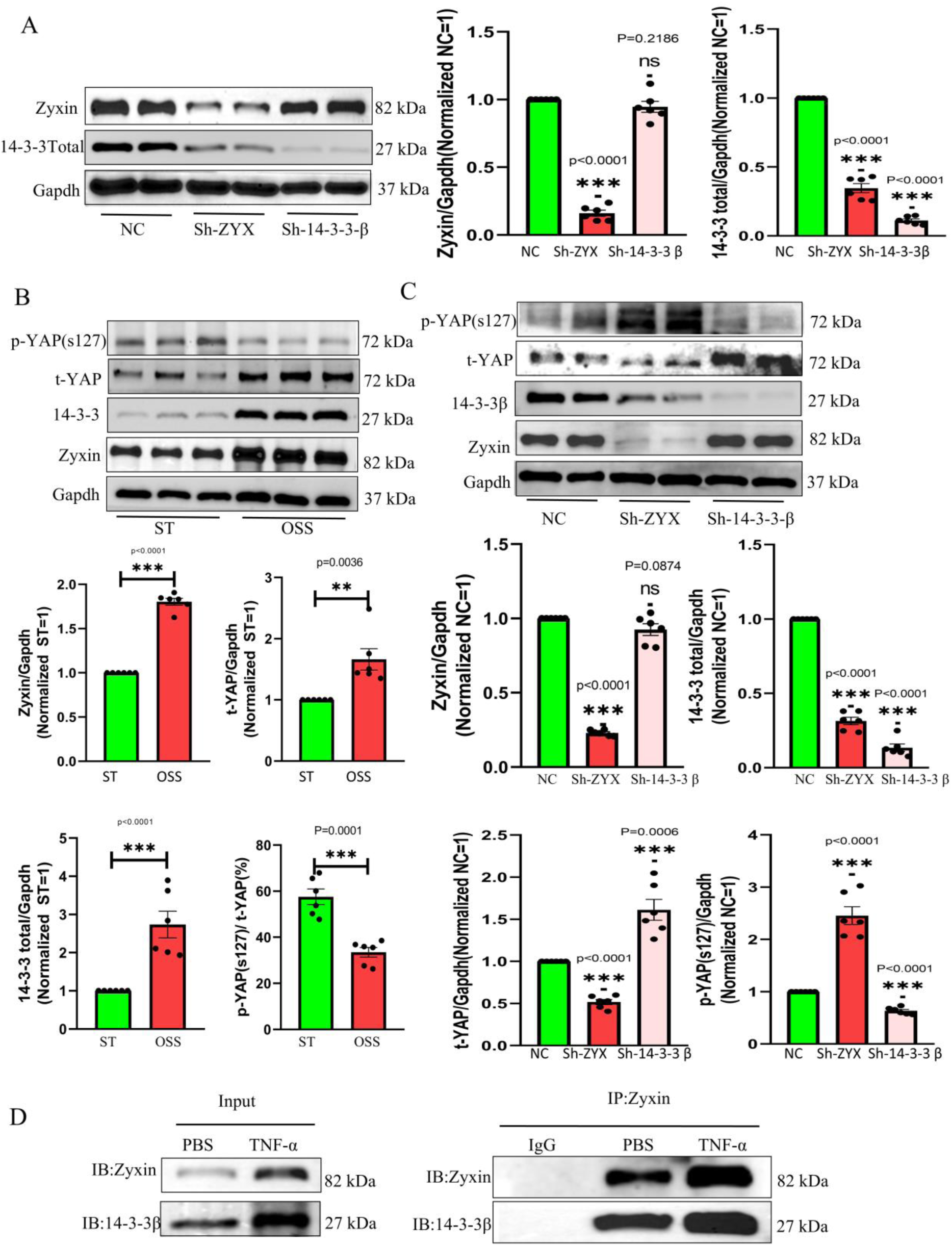
Zyxin-14-3-3 axis mediates YAP nuclear translocation to promote endothelial inflammation. **A,** Immunoblotting results showed that loss of function of zyxin in HUVECs significantly down-regulated the level of total 14-3-3 protein expression compared with the NC group. Whereas the loss of function of 14-3-3β (YWHAB), the protein expression of Zyxin in HUVECs was not statistically different compared with the NC group. ST, static; OSS, oscillatory shear stress. n=6. **B-C,** Immunoblotting results showed that YAP phosphorylation at the S127 site was inhibited and YAP expression was up-regulated when HUVECs were exposed to OSS for 6 h (B), ST, static; OSS, oscillatory shear stress. Zyxin loss of function in HUVECs promoted phosphorylation of YAP at the serine S127 site, and YAP protein expression was inhibited. Upon loss of function of 14-3-3β, YAP phosphorylation at the serine S127 site was inhibited, promoting YAP protein expression (C), n=6. **D,** Co-immunoprecipitation of zyxin and 14-3-3β in HUVECs treated with TNF-alpha (10 ng/mL) for 6 hours. n=6. A-C Statistical analysis was performed by Mann-Whitney test.

### 7. Rosuvastatin suppresses zyxin expression and protects against atherosclerosis

To determine whether existing anti-atherosclerotic drugs inhibit zyxin expression, the clinically utilized drug Rosuvastatin was investigated. It was confirmed that treatment with Rosuvastatin significantly reduced zyxin gene expression in HUVECs (Figure S8A). Furthermore, the OSS-stimulated expression of both zyxin and YAP was markedly diminished, and the OSS-induced expression of VCAM-1 was mitigated (Figure S8B). Additionally, Rosuvastatin suppressed the mRNA levels of YAP downstream target genes CTGF and Cyr61, as well as the inflammatory factors ICAM-1 and MCP-1 (Figure S8C-F). To further validate the effect of Rosuvastatin on zyxin, zyxin was knocked down in the medium, and HUVECs were subsequently treated under either static conditions or OSS for 6 hours following a 30-minute pretreatment with PBS or Rosuvastatin. The expression levels of zyxin and VCAM-1 were assessed. Results indicated that Rosuvastatin effectively suppressed OSS-induced VCAM-1 expression, while the knockdown of zyxin prevented the elevated VCAM-1 protein levels induced by OSS. Notably, no statistically significant differences were observed in the protein expression levels of zyxin and VCAM-1 between the Rosuvastatin-treated group and the control group after zyxin deletion (Figure 7A). These findings suggest that the downregulation of zyxin expression in ECs may represent one mechanism through which Rosuvastatin suppresses OSS-induced ECs inflammation. Subsequently, the effect of Rosuvastatin on zyxin expression in vivo within the endothelium predisposed to AS was examined. Eight-week-old male ApoE^-/-^ mice were subjected to either sham surgery or partial ligation of the left common carotid artery to establish a mouse model of OSS-induced AS. Two weeks later, during the treatment phase, the sham operation group and the PLCA group were administered either 10 mg/kg of Rosuvastatin or an equivalent volume of PBS (Figure 7B). At the conclusion of the treatment, aortic tissues were collected from each group, and the plaques in the left common carotid artery were assessed. Oil Red O staining revealed that Rosuvastatin significantly decreased the area of atherosclerotic plaques in PLCA ApoE^-/-^ mice (Figure 7C-D). Western blot analysis indicated that the protein expression of endothelial zyxin and VCAM-1 was markedly reduced in the Rosuvastatin treatment group compared to the surgery group (Figure 7E). RT-qPCR analysis demonstrated that Rosuvastatin inhibited the expression of zyxin, YAP, and its target genes CTGF and Cyr61, as well as the inflammatory factors VCAM-1, ICAM-1, and MCP-1 at the mRNA level in the PLCA vascular tissues (Figure S8G). These results suggest that Rosuvastatin suppresses zyxin expression and mitigates the expression of YAP and inflammatory factors, exerting both anti-inflammatory and anti-atherosclerotic effects. This indicates that zyxin may hold potential as a therapeutic target for AS.

**Figure 7.**
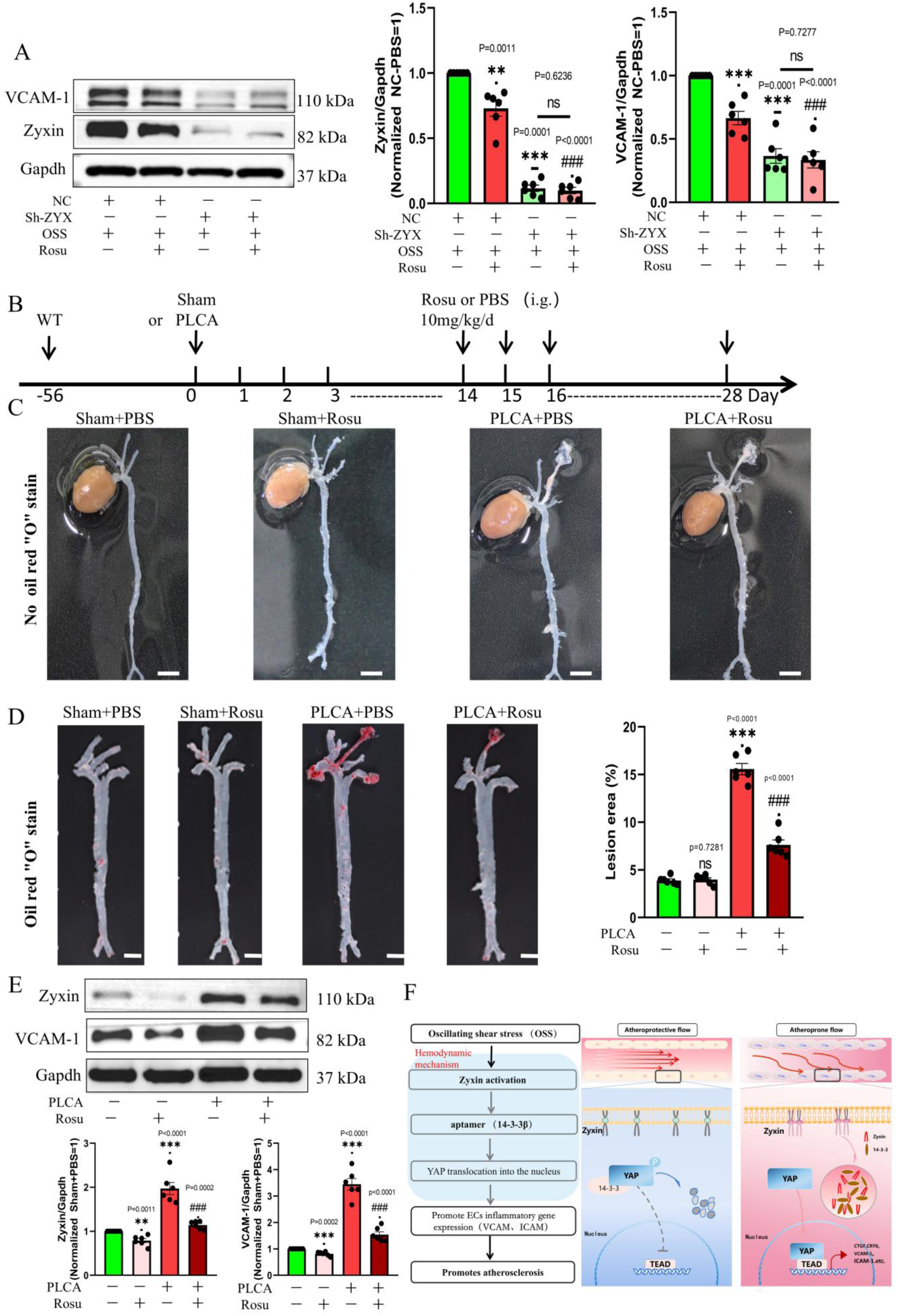
Rosuvastatin Suppresses Zyxin Expression and Protects Against Atherosclerosis. **A,** Western blotting showed that rosuvastatin treatment reduced the expression level of zyxin and VCAM-1 in HUVECs exposed to OSS, while there was no significant difference in the protein level of VCAM-1 between the rosuvastatin treatment group and the control group following zyxin loss-of-function. N=6, OSS, oscillatory shear stress, Resu, Rosuvastatin. **B,** Rosuvastatin administration schedule in ApoE^-/-^ mice treated with sham surgery or PLCA. PLCA, Partial ligation of left common carotid artery; Resu, Rosuvastatin. C-D, Aortic artery tissue was obtained from ApoE^-/-^ mice four weeks following partial ligation of the left common carotid artery for the purpose of examining atherosclerotic lesions in the control and rosuvastatin groups. **C,** No oil red staining; scale bar:1mm, n=6. **D,** Arterial tissues were used for oil red staining and quantification of the lesion area. Scale bar: 1mm, n=6. PLCA, Partial ligation of left common carotid artery; Resu, Rosuvastatin. **E,** Immunoblotting showing zyxin and VCAM-1 protein levels in the left common carotid artery after sham operation or ligation of PBS and rosuvastatin treated mice. Quantitative data are shown in the figure ring, n=6. PLCA, Partial ligation of left common carotid artery; Resu, Rosuvastatin. **F,** Schematic overview of endothelial zyxin regulation and signaling in blood flow-associated atherosclerosis. Statistical analysis was performed by 2-way ANOVA followed by the Tukey test for A, D and E.

## DISCUSSION

Atherosclerosis (AS) remains a leading cause of cardiovascular disease (CVD) worldwide, serving as a common pathological basis for several critical conditions, including coronary heart disease, stroke and aortic dissection, which pose a significant threat to human health^1^. Extensive studies have demonstrated that AS is not merely a disorder of lipid metabolism; conversely, inflammation plays a vital role in both the initiation and the advancement of its complications, exerting an influence on the disease as a whole^3,5,9,34^. Despite recent clinical advancements in anti-inflammatory therapies targeting molecules such as IL-1β and NLRP3^35,36^, the prevalence of AS and its associated vascular diseases remains unchanged^1,2^. Therefore, it is imperative to investigate upstream pathological factors and identify key therapeutic molecules involved in the progression of AS^1,2,37,38^. This study emphasizes hemodynamics, utilizing ECs as the primary research subjects and employing inflammation as a guiding framework. The objective is to explore the early initiating factors of AS and to identify effective therapeutic targets.

This study demonstrates that zyxin responds to flow shear stress and acts as a crucial regulator of ECs inflammation. Its response to OSS promotes the progression of AS, as illustrated in Figure 7F. This conclusion is supported by subsequent in vivo and in vitro findings: (I) Zyxin mRNA and protein levels are significantly elevated in arterial plaques from patients with coronary heart disease. The deletion or knockdown of zyxin results in a marked reduction in the expression of endothelial adhesion molecules and pro-inflammatory cytokines in ECs exposed to OSS, thereby mitigating the progression of AS induced by turbulent blood flow or a Western diet in mice. (II) Under OSS or TNF-α stimulation, zyxin mediates the nuclear translocation of YAP. Pre-treatment of ECs with the non-specific YAP inhibitor verteporfin counteracts the pro-atherosclerotic effects mediated by zyxin, indicating that Hippo-YAP signaling is involved in zyxin-mediated AS. (III) Zyxin does not interact directly with YAP but regulates 14-3-3β to inhibit YAP phosphorylation at serine 127, subsequently promoting YAP expression and nuclear translocation, which enhances endothelial inflammation and the expression of genes related to adhesion factors involved in AS pathogenesis. (IV) The expression of zyxin in the vascular endothelium in regions susceptible to AS is downregulated by Rosuvastatin, alleviating zyxin-mediated endothelial inflammation and AS. These results suggest that inhibiting the activation of the zyxin-14-3-3β-YAP signaling axis could represent a promising anti-inflammatory strategy for treating AS. Furthermore, these findings reveal previously unknown functions of zyxin, positioning it as a key molecular determinant of mechanical forces, cytoplasmic signaling cascades, and intercellular communication within the extracellular environment. This unified theoretical framework elucidates the molecular mechanisms underlying ECs activation induced by OSS.

ECs possess numerous mechanosensory microstructures that respond to mechanical stimuli, enabling them to sense and transduce biomechanical signals into biochemical responses that regulate homeostatic and vascular functions^5,6,12,39^. These sensors and microstructural domains include a variety of proteins and molecules, such as cell-cell junction proteins, adhesion molecules, receptors, ion channels, membrane microdomains, and glycocalyxes ^12,39^. Focal adhesions are large multiprotein complexes within the extracellular matrix, consisting of over 150 proteins, that transduce mechanical signals into the cell^12,16,40^. Zyxin functions as a mechanotransduction molecule^12,19,21,41,42^, and previous studies examining zyxin’s response to mechanical stimuli have primarily focused on mechanical stiffness and stretch^20,22,42,43^. For instance, cyclic stretch has been shown to induce the transfer of zyxin from focal adhesions to the actin cytoskeleton^20,40–42^. In the realm of disease research, studies involving zyxin-related diseases have predominantly centered on tumors, including those of the breast and colon, as well as tissue repair^15,20,30,42,44^. Recently, there has been a notable increase in research on zyxin and cardiovascular diseases, particularly concerning hypertension and cardiovascular embryonic development^21,23,45,46^. This study demonstrates that zyxin, a mechanosensitive transduction molecule, responds to shear stress and is linked to ECs inflammation. Elevated zyxin expression in ECs promotes inflammation and exacerbates AS, while the deletion of zyxin inhibits inflammation and alleviates atherosclerotic lesions. These findings enhance our understanding of the role of zyxin.

Flow shear stress is transmitted from the plasma membrane to the nucleus of ECs via mechanoreceptors, junctional proteins, kinases, and transcription factors, which collaborate at various levels to control the expression of genes that determine cell fate and phenotype^4,5,7,12,39^. In this study, we found that zyxin may mediate OSS-induced ECs inflammation by regulating YAP, a key molecule in the Hippo signaling pathway, through transcriptomic analysis and exploration of a protein interaction network (STRING). The Hippo signaling pathway is a highly conserved pathway that has gained attention in recent years^24,25,29,47^. In mammals, it includes several key components: MST1/2, SAV1, MOB1A/B, LATS1/2, YAP1, TAZ, and TEAD^24,27,31,48^. YAP and TAZ act as transcriptional co-activators, binding to TEAD1-4, which are crucial mediators of mechanical signaling regulation^26,49–51^. Dysregulation of the Hippo pathway is linked to various diseases, including cancer, eye disease, heart disease, lung disease, kidney disease, liver disease, and immune dysfunction^24,26,51^. The activation of the Hippo pathway restrains the activity of YAP/TAZ through the phosphorylation mediated by LATS1/2^27,30,50,51^. Conversely, The deactivation of the Hippo pathway brings about the translocation of dephosphorylated YAP/TAZ to the nucleus and its binding to TEAD1-4, which leads to the induction of gene expression^25,31,47,51^. Previous studies have shown that OSS promotes nuclear translocation of YAP, a central molecule in the Hippo pathway, and is closely associated with EC inflammation and AS^5,25,29,47,50^. However, the mechanisms governing YAP nuclear translocation remain poorly understood. Some studies indicate that β-linker protein complexes with YAP and TBX5, inducing the transcription of anti-apoptotic genes post nuclear translocation^52^; BACH1 forms a transcription factor complex with YAP, contributing to the activation of inflammatory genes via YAP nuclear translocation^53^. Our study suggests that zyxin does not directly regulate YAP; rather, it influences YAP nuclear translocation by interacting with 14-3-3β to depolymerize the 14-3-3-YAP complex. The 14-3-3 proteins are a highly conserved family of proteins with seven isoforms in mammals: β, γ, ε, ζ, η, τ, and σ, which collectively function as regulatory proteins^54^. It is established that 14-3-3 proteins act as adaptor proteins for over 200 target proteins^28,54,55^, and there is growing evidence linking changes in 14-3-3 protein levels to chronic inflammatory diseases^28,54,55^. This study showed that 14-3-3β is the predominant isoform of the 14-3-3 protein family within HUVECs, suggesting that zyxin acts as an upstream regulator of 14-3-3β. Zyxin was shown to interact with the 14-3-3β protein via co-immunoprecipitation (CO-IP). These findings suggest that zyxin may play a role in regulating OSS-induced YAP nuclear translocation to promote ECs inflammation by modulating 14-3-3β.

Clinical studies, such as CANTOS, along with various basic cell biology research, have established that inflammation is recognized as a key driver of AS^3,35,37,38^. It has been shown that statins exert their anti-atherosclerotic effects primarily by reducing lipid levels and inflammation^35,37,38^. The cytoskeletal protein zyxin is involved in numerous cellular functions^18,21,22,32,41,45,56^. In the present experiments, it was found that zyxin responds to OSS to promote ECs inflammation and participate in the pathogenesis of AS. A significant alleviation of OSS-induced ECs inflammation and a reduction in the area of atherosclerotic lesions were observed with zyxin deficiency. This suggests that targeting zyxin may represent a valid therapeutic strategy for combating both AS and inflammation. The investigation into whether the anti-inflammatory and anti-atherosclerotic properties of Rosuvastatin are associated with zyxin was also conducted. The experimental results indicated that a significant reduction in the expression levels of zyxin and YAP was achieved with Rosuvastatin, which inhibited ECs inflammation and diminished the area of atherosclerotic plaques in partially ligated left carotid arteries of APoE^-/-^ mice exposed to OSS. These findings suggest that zyxin may play a role in the anti-atherosclerotic effects of Rosuvastatin. Previous studies have demonstrated that the anti-atherosclerotic effects of Rosuvastatin are mediated by inhibiting zyxin expression, indicating that stabilizing cytoskeletal proteins could be an effective strategy for treating AS and holds promise for future therapies. ECs rapidly sense changes in flow shear stress, which influences the progression of atherosclerotic plaques. This study demonstrates that zyxin acts as a mechanosensor for shear stress, mediating endothelial inflammation and AS induced by OSS. The 14-3-3β protein is identified as the primary isoform of the 14-3-3 protein family involved in the response to OSS, receiving regulatory signals from zyxin and mediating the nuclear translocation of YAP. Overall, we elucidate that the zyxin-14-3-3β-YAP axis plays a crucial role in endothelial inflammation and AS. Furthermore, these findings may enhance our understanding of how the Hippo pathway regulates and contributes to the development and progression of AS.

## Supplemental Material

Expanded Materials and Methods

Figures S 1–S 8

Tables S 1 and S 5

**Reference** 26, 31, 42

